# A Quantitative Measurement to Describe the Relative Proximity of Fluorescent Biomarkers

**DOI:** 10.1101/2020.12.08.416206

**Authors:** Joshua D. Dyer, Alan R. Brown, Alex Owen, Jeremy Metz

## Abstract

Determining the relationship between biomarkers via fluorescence microscopy is a key step in the characterisation of cellular phenotypes. We define a simple distance-based measurement termed a perimeter distance mean (PD_mean_) which quantifies the relative proximity of objects in one fluorescent channel to objects in a second fluorescent channel in 2D or 3D microscopy datasets. PD_mean_ measurements were able to accurately identify known changes in colocalisation in computer-generated and real-world microscopy datasets. We argue that this approach provides substantial advantages over currently used distance-based colocalisation analysis methods. We also introduce PyBioProx, an extensible open-source Python module and graphical user interface that produces PD_mean_ measurements.

## Introduction

Determining the colocalisation of fluorescent biomarkers via microscopy is a frequently employed method to infer meaningful biological interactions within cells ^1^. Researchers take single or multiple images (Z-stack) along the Z-axis in the same X/Y axes to record the distribution of fluorescent biomarkers in 2D or 3D space. Analysis methods varying in complexity are then employed to quantify the relationship between the biomarkers ^2^. A commonly employed semi-quantitative analysis involves overlaying pseudo-labelled (e.g. red and green) fluorescent channels and manually counting regions of (yellow) overlap as instances of biomarker colocalisation. This method, however, is time-consuming and highly subjective. Detection of (yellow) regions of overlap only occurs when histograms of each channel are similar, likely not to be the case when imaging fluorochromes with different signal strengths ^2^. Therefore, erroneous identification of colocalisation may occur as overlap identification depends both on the calibration of the display equipment and the perceptions of the investigator ^3^.

Quantitative colocalisation analysis techniques can be broadly grouped into pixel-based ^3,4^ and object-based methods ^5–7^, alongside methods that use a combination of both approaches ^8,9^. Strictly pixel-based analysis methods use statistical approaches to determine the degree of overlap in fluorescent signals globally (i.e. throughout the image on a pixel-by-pixel basis) ^5^. Object-based methods, by contrast, require that fluorescent objects first be identified and separated from surrounding parts of the image. Spatial information included in the metadata of the microscopy image is then used to define the relationship between the objects ^7^.

In this paper, we describe a simple object-based measurement that quantifies the relative proximity of objects in one fluorescent channel to objects in a second fluorescent channel. This method can identify changes in the spatial proximity of fluorescently-labelled biomarkers that occur according to experimental conditions. For example, one could use this method to investigate whether the relative proximity of fluorescently-labelled proteins or organelles (e.g. the proximity of mitochondria to the endoplasmic reticulum) changes following exposure of a cell to a particular treatment. We use this approach to quantify how the spatial proximity of intracellular *Staphylococcus aureus* relative to the lysosomal-associated membrane protein 1 (LAMP-1), changes over time.

Perimeter Distance (PD) measurements are defined in Fig 1 and are detailed extensively in the methods section. In brief, objects in one fluorescent channel are detected, and the perimeter pixels around the object determined. The distance of each pixel in an object’s perimeter to the nearest detected object in the second channel is then calculated. The units of PD measurements may be pixels, voxels or real-world units (e.g. µm/nm) if pixel/voxel dimensions are known. Depending on the number of perimeter pixels, each object may have thousands of PD measurements. We assessed the capacity of the mean (PD_mean_), minimum (PD_min_) and maximum (Hausdorff Distance) PD measurement to accurately identify changes in colocalisation. Across both computer-generated and real-world datasets, we found that the PD_mean_ measurement (but not Hausdorff Distance or PD_min_ measurements) was consistently able to identify known changes in colocalisation.

**Figure 1.**
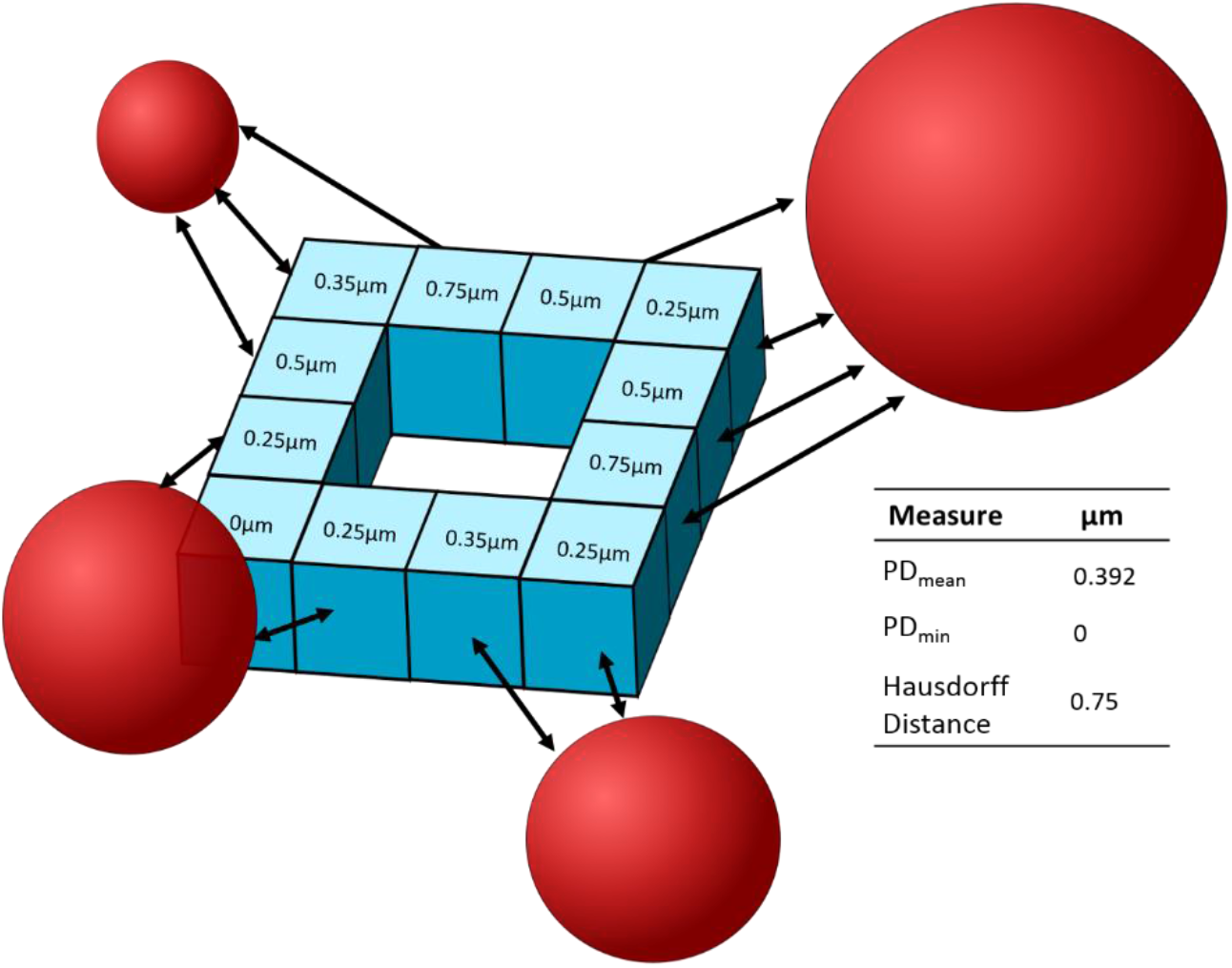
PD Measurements. An illustrative example in which the proximity of a blue fluorescent object relative to red fluorescent objects is assessed. 12 perimeter voxels (illustrated as light-blue boxes) have been identified for the blue object. The numbers in the centre of each perimeter-voxel refer to PD measurements: the shortest distance from the perimeter-voxel to the nearest object in the red fluorescent signal. The mean of these PD measurements gives the PD_mean_. The largest and smallest of these PD measurements gives the Hausdorff Distance and PD_min_ distances respectively.

Analysis of biomarker association by distance measurements is not a novel concept. For example, the ‘nearest-neighbour distance’ analysis proposed by Lachmanovich and colleagues (2003) considers two objects as colocalised if their centroids are within a threshold distance ^6^. However, the binary nature of this analysis (where objects are either colocalised or not), results in the loss of large amounts of data. An approach described by several groups and implemented in the DiAna ImageJ plugin produces continuous data by collecting measurements between pairs of differently-labelled fluorescent objects (e.g. by measuring the distance between the centroids of objects (centroid-centroid measurements) ^5,7,10,11^. In contrast to the methods described above which measure the distances between *pairs* of differently-labelled fluorescent objects, the PD_mean_ measurements described in this paper produce continuous data describing the position of a fluorescent object in one channel relative to *all* surrounding fluorescent signal in another fluorescent channel. We argue herein that this provides a more robust measure of the relative proximity of one fluorescent biomarker to another.

We also introduce PyBioProx, an open-source tool written in the Python programming language that generates PD measurements to describe the relative proximity of fluorescent biomarkers in 2D or 3D space. For users familiar with Python, PyBioProx is available as an extensible Python module. For users unfamiliar with Python, PyBioProx is available as a user-friendly graphical user interface (GUI) that can be run from the command line. Other core advantages that PyBioProx brings include a high aptitude for the batch distance-based analysis of large 2D and 3D datasets, as well as the extensibility of being written in an easy-to-learn and fast-to-prototype programming language. Detailed instructions for PyBioProx installation and use are available at **pybioprox.github.io**

## Methods

### PyBioProx Distance Analysis

PyBioProx computes distances between labelled objects in separate fluorescence channels. Before the detection of fluorescent objects, different pre-processing steps can be performed to improve detection accuracy. Due to its extensible nature, user-selected pre-processing functions can easily be passed into the detection function using the straight-forward module application programming interface (API). A second alternative is to pre-process images with another program (e.g. ImageJ) and pass in pre-processed images directly into PyBioProx. Images analysed in Figs 3 & 4 were pre-processed with an unsharp-mask in the LAMP-1 channel in ImageJ (sigma = 10, mask weight = 0.6) followed by a three-dimensional Gaussian blur in both channels (sigma = 1) performed in Python using the scipy.ndimage.gaussian_filter function. No pre-processing was employed on the images analysed in Figs 2 and 5.

**Figure 2.**
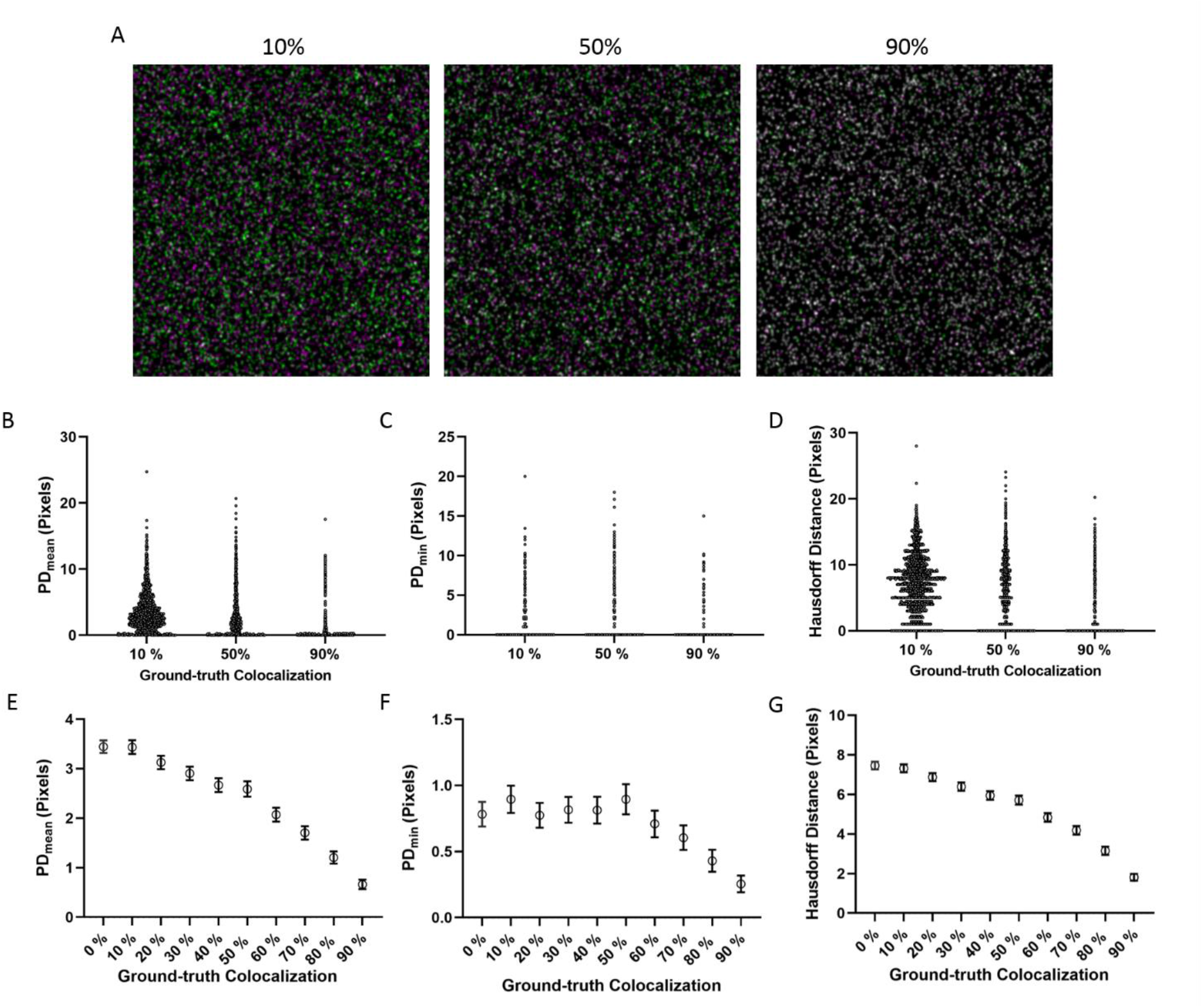
PD measurements identify known changes in colocalization. (A) Representative images of CBS ‘Image Set 1’ with known ground-truth colocalization values. The ‘red’ channel in the CBS ‘Image Set 1’ is displayed in magenta for better visualization. (B-D) PD_mean_/PD_min_ /Hausdorff Distance values for each of the magenta objects relative to green objects in (A) calculated using PyBioProx. (E-G) The PD_mean_/PD_min_ /Hausdorff Distance values for objects in the magenta channel of the entire CBS dataset 1. Values represent the mean ∓ 95 % CI for each image.

**Figure 3.**
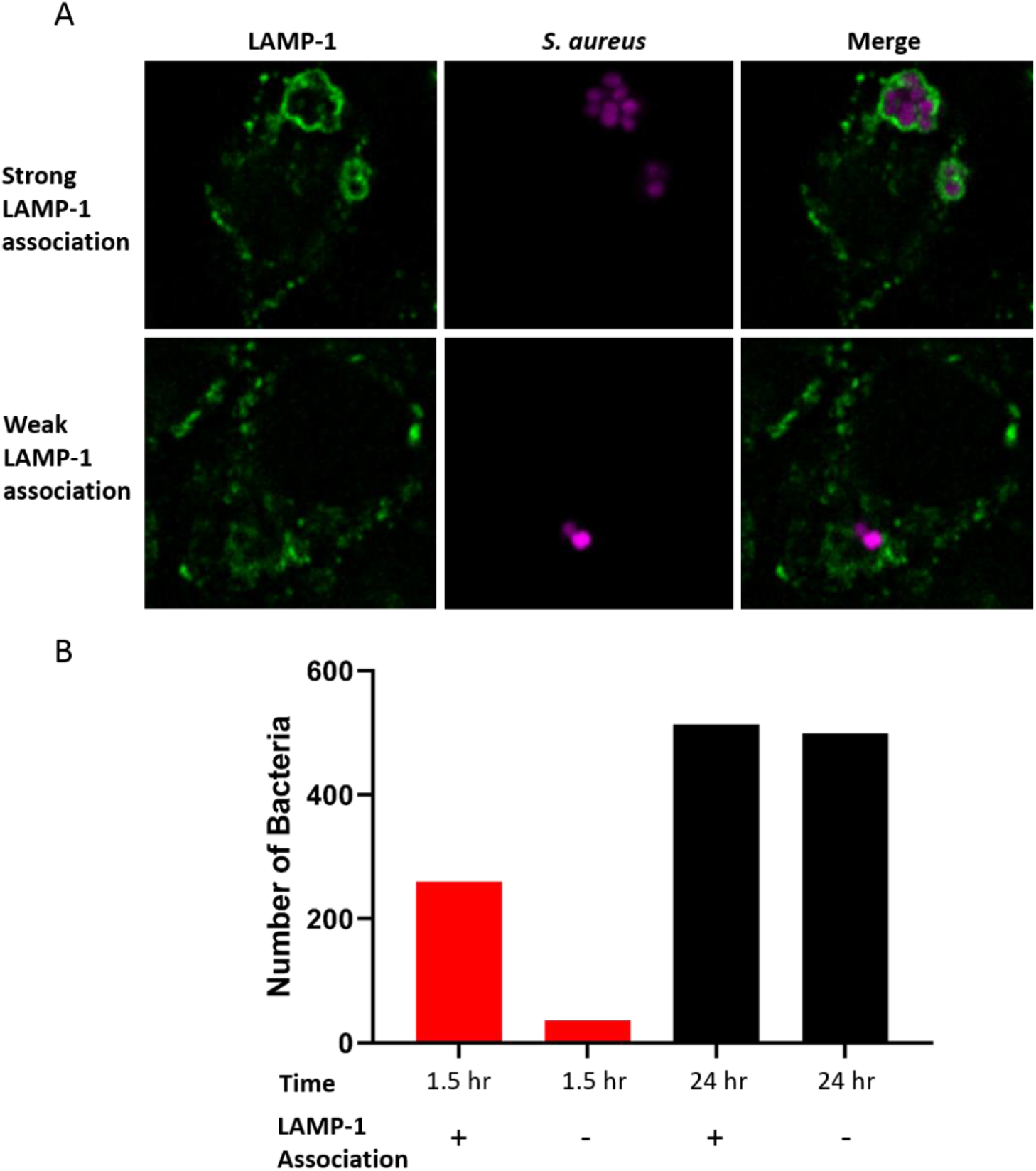
The association between *S. aureus* MSSA476 and LAMP-1 in J774 macrophages changes over time. (A) Example of weakly and strongly LAMP-1-associated *S. aureus*. A single slice of a Z-stack is shown. (B) Manual analysis of *S. aureus*-LAMP-1 association. Images were blinded and the number of *S. aureus* with/without halos of LAMP-1 signal around them were counted as positive (+) and negative (−) LAMP1-associated bacteria respectively.

Following pre-processing, PyBioProx generates binary 2D/3D masks to define regions containing foreground or background pixels, using binary values ‘True’ (1) and ‘False’ (0) to represent presence or absence of signal. The PyBioProx GUI provides a range of thresholding methods from the scikit-image package to define what is selected as positive and negative signal ^12^. The Otsu thresholding method is used in the examples presented in this paper ^13^. A comparison operator then generates binary 2D/3D images. Alternatively, users can provide PyBioProx with binary images previously thresholded in another program. Connected regions of ‘on’ pixels are then labelled as unique objects using the scipy.ndimage.label function. A binary erosion of the labelled objects is then employed to identify object perimeter pixels. Finally, overlays of the detected object’s perimeter on (user-defined) representative Z-slices of the original image are generated to allow for confirmation that fluorescent objects have been appropriately labelled.

### Macrophage Infection Assay

J774A.1 cells were routinely cultured in DMEM containing 4500 mg/L glucose and 10 % FBS and incubated at 37 °C with 5 % CO_2_. Three replicate wells were seeded with 2.5 × 10^5^ cells in 24 well plates containing glass coverslips and grown overnight. Cells were washed 2x with PBS before inoculating with *S. aureus* MSSA476 carrying the pRN11 (mCherry) expression plasmid at an MOI of 1. Infection was allowed to proceed for 2 hrs, before washing with PBS and treating with 100 µg/ml gentamicin in DMEM for 1 hr to kill extracellular bacteria. Cells were then washed 3x with PBS and DMEM containing 10µg/ml gentamicin added to prevent bacterial outgrowth. At relevant time points, cells were washed with PBS and fixed with 4% paraformaldehyde for 10 min at 37°C. Cells were washed with PBS and blocked/permeabilised in a PBS solution containing 0.2 % saponin & 5 % bovine serum albumin (BSA). Cells were washed in PBS, overlaid with primary antibody (1:100 in blocking/permeabilisation solution) and incubated overnight at 4°C. Cells were extensively washed in PBS then overlaid with secondary antibody (1:400 in blocking/permeabilisation solution) for 1 hr before washing in PBS and mounting on slides containing prolong gold. Mounted samples were imaged using a Zeiss LSM 880 laser scanning microscope using the Alpha Plan Apochromat 100x/1.46 (NA) oil DIC M27 objective. LAMP-1 fluorescence was excited with an Argon laser at 488 nm and detected between 493-579 nm (pinhole size = 1 AU). S. aureus-mCherry fluorescence was excited using an He 543 laser at 561 nm and detected between 579-696 nm (pinhole size = 0.84 AU). Voxel scaling = 0.08 μm × 0.08 μm × 0.75μm (XYZ). 5 Z-stack images were taken per technical replica.

## Results & Discussion

### 1. PD_mean_ measurements correctly identify known differences in a pixel-based colocalisation dataset

The Colocalization Benchmark Source (CBS) is an online database of 2D computer-simulated images with pre-defined (ground-truth) values of pixel-based colocalisation ranging from 0-90 % ^14^. To validate the capacity of this approach to detect changes in pixel-level colocalisation, we utilised ‘Image Set 1’ from the CBS. Images were processed using PyBioProx as described in the methods section. Example CBS images with ground-truth colocalisation values of 10%, 50% and 90% are shown (Fig 2A). The PD_mean_ / PD_min_ / Hausdorff Distance for each magenta object relative to green objects was calculated (Fig 2B-D) and the distributions of the 10 %, 50 % and 90 % images displayed as a scatter plot (Fig 2E-G). 1544, 1592 and 1588 magenta objects were detected for the 10 %, 50 % and 90 % images respectively. As the ground-truth colocalisation value of the images increases, the PD_mean_ and Hausdorff Distance values for the magenta objects within these images cluster progressively closer to 0 pixels (Fig 2B & D). The PD_min_ values (Fig 2C) of the objects do not show a similar pattern.

Across the entire CBS dataset (CBS ‘Image Set 1’ (Fig 2E & G), ‘Image Set 2’ and ‘Image Set 3’ (Supp. Fig 1A - D)), reductions in PD_mean_ and Hausdorff Distance values are observed with increasing ground-truth colocalisation. While the changes in PD_mean_ and Hausdorff distance values relative to ground-truth colocalisation are not as linear as can be obtained using pixel-based colocalisation metrics (e.g. the protein proximity coefficient ^14^), it is impressive that object-based measurements such as the PD_mean_ and Hausdorff Distance, are able to identify changing colocalisation values in a pixel-based colocalisation dataset. By contrast, PD_min_ values do not consistently decrease with increasing ground-truth % colocalisation (Fig 2C & F, Supp Fig 1E & F). This poor performance reflects that only the closest pixel in a magenta object to green fluorescent signal is used in PD_min_ measurements. As such, a poorly colocalising object with a PD_mean_ distance of 10 pixels and a Hausdorff Distance of 20 pixels, could conceivably have a PD_min_ distance of 0 pixels. As the CBS datasets represent crowded images, with large numbers of objects in close proximity to one-another, PD_min_ distances are inappropriate and likely to give erroneous results.

### 2. PD_mean_ measurements correctly identify changes in biomarker proximity in a real-world dataset

The internalisation of bacteria by innate immune cells such as macrophages is an important mechanism to contain microbial threats. Phagocytosis is a complex process involving the internalisation of foreign particles such as bacteria and apoptotic cells into a benign compartment termed the nascent phagosome ^15^. A complex series of maturation events then occur in which the nascent phagosome rapidly changes its membrane composition and inter-luminal contents to form a microbicidal compartment termed the phagolysosome ^16^.

Lysosome-Associated-Membrane-Protein 1 (LAMP-1) is a regularly utilised biomarker of the phagolysosome ^17–21^. The capacity of *S. aureus* to survive for extended periods of time within macrophages has become increasingly apparent ^17–19,22,23^. Recent work indicates a capacity of *S. aureus* to survive and even replicate within LAMP-1 positive vesicles ^17,18^. Escape of *S. aureus* strains from the phagolysosomes of primary human macrophages and THP-1 cells has also been reported ^17,22^. We prepared an infection model of *S. aureus* MSSA476 in J774A.1 murine macrophage-like cells, with LAMP-1 staining. At early (1.5 hr) and late (24 hr) timepoints, Z-stack images were captured. Significant overlap in the fluorescent signal is not necessarily expected between bacteria and LAMP-1. Instead, a ‘halo’ of LAMP-1 around a bacterium (Fig 3A) indicates the localisation of the bacteria within a phagolysosome ^17–21^.

Typically, analyses of the extent of LAMP-1 encapsulation around bacteria are user-defined binary measures; manually counting the number of positive/negatively LAMP-1-associated bacteria in a dataset of blinded images ^17–21^. This method of analysis is time-consuming and raises issues of reproducibility. Defining what amount of LAMP-1 association equates to a positively LAMP-1 associated bacterium is subjective and may vary from researcher to researcher. This subjectivity may lead to overselling or underselling results based on the definition of ‘positive association’. The binary nature of the analysis may also result in the loss of important information as small but meaningful changes in the extent of bacterial encapsulation by LAMP-1 may not be identifiable by eye. Fig 3B shows the results of a manual analysis in which images were blinded and the number of positively/negatively encapsulated bacteria counted in Image J. At the 1.5 hr time point, 87.8 % of bacteria were counted as LAMP-1 associated. At the 1.5 hr time point, 87.8 % of bacteria were counted as LAMP-1 associated, compared to only 50.7 % at the 24 hr time point, potentially indicating the escape of *S. aureus* from LAMP-1 positive phagosomes. Notably, a 3.4 fold increase in the number of bacteria detected at 24 hr (1012 bacteria) compared to 1.5 hrs (296) was observed.

The same dataset was then analysed using PyBioProx. Prior to both the manual and PyBioProx analysis, the LAMP-1 channel was pre-processed by an unsharp mask in ImageJ. Unsharp masking produces a sharpened image by subtracting a blurred copy of the image from the original and rescaling the histogram to produce the original contrast in low-frequency features ^24^. Unsharp masking resulted in a more discrete detection of positive LAMP-1 signal (Supp. Fig 2A) and a greater capacity to resolve differences in colocalisation as evidenced by larger PD_mean_ and Hausdorff Distance values (Supp. Fig 2B). Further pre-processing was performed using a Gaussian filter (scipy.ndimage.gaussian_filter) to reduce single-pixel scale noise. In the absence of using the Gaussian filter, large numbers of single-pixel *‘S. aureus* objects’ were erroneously identified. Following pre-processing, the PD_mean_, Hausdorff Distance and PD_min_ for each bacterial cluster in 3D space at early and late time points were calculated in PyBioProx relative to LAMP-1 fluorescence (Fig 4A, C & E). A reduced number of *S. aureus* objects were identified by PyBioProx (495) compared to the number identified by manual analysis (1308). This reflects the limited capacity of the thresholding methods used, to segment large clumps *of S. aureus* into individual cells. Consequently, *S. aureus* objects detected by PyBioProx are referred to as ‘*S. aureus* clusters’. In this dataset, objects composed of fewer than ten connected pixels are considered to be noise and removed from the analysis. The mean *S. aureus* object in the dataset contained 866.6 ± 84.72 (SEM) perimeter pixels. As in the manual analysis (Fig 3B), a larger number of bacterial clusters were detected by PyBioProx at the 24 hr time point (395) then at the 1.5 hr time point (100).

**Figure 4.**
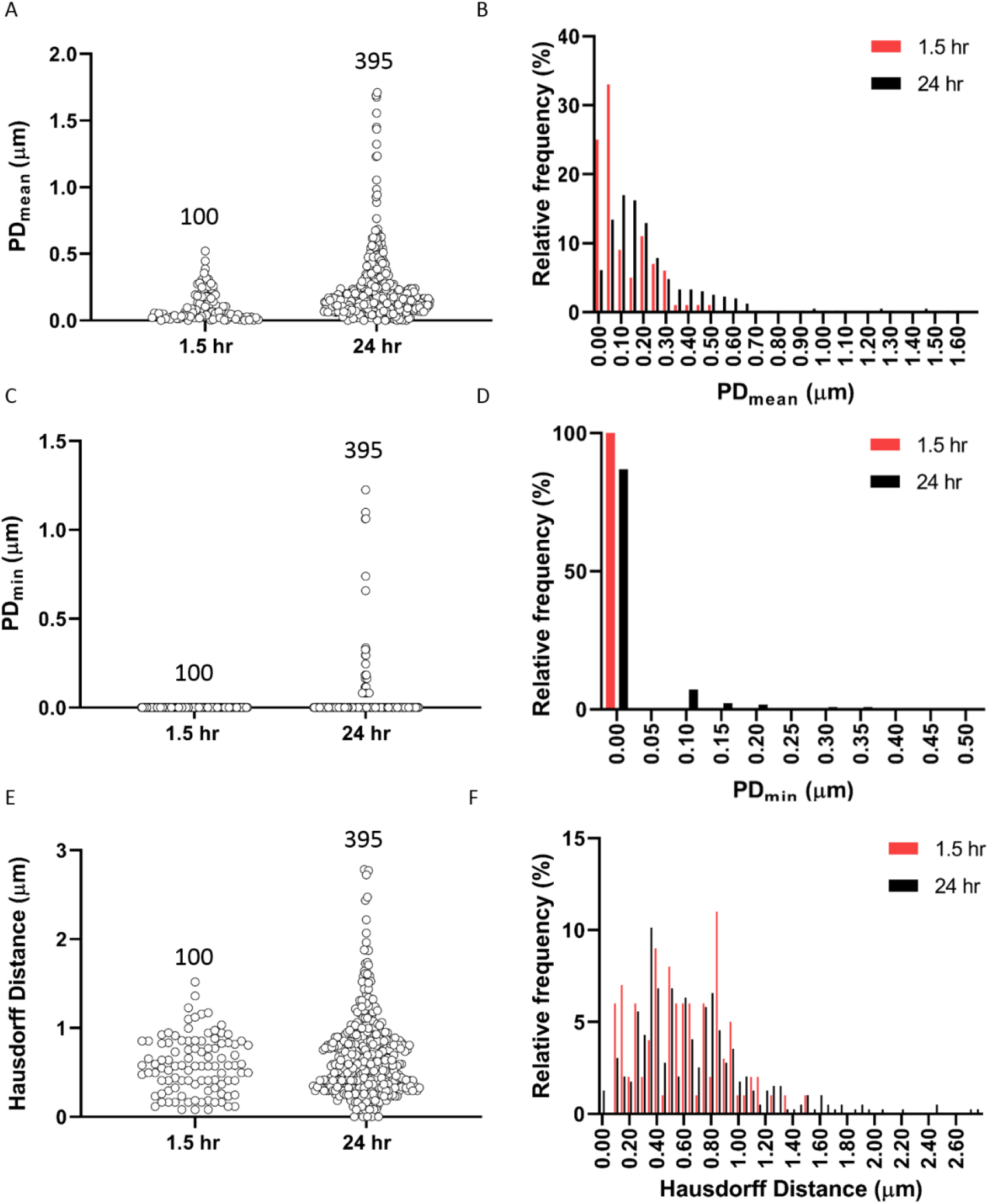
PyBioProx Analysis of LAMP-1-*S. aureus* association. (A, C & E) PD_mean_/Hausdorff Distance/PD_min_ measurements describing the relative proximity of *S. aureus* to LAMP-1. Each dot represents the PD_mean_/Hausdorff Distance/PD_min_ distance value for an individual bacterial cluster. The number of bacterial clusters identified is displayed on top of each timepoint. (B, D & F) Frequency distributions of the *S. aureus* PD_mean_/Hausdorff Distance/PD_min_ values (in A, C & E respectively) at the 1.5 hr and 24 hr timepoints.

**Figure 5.**
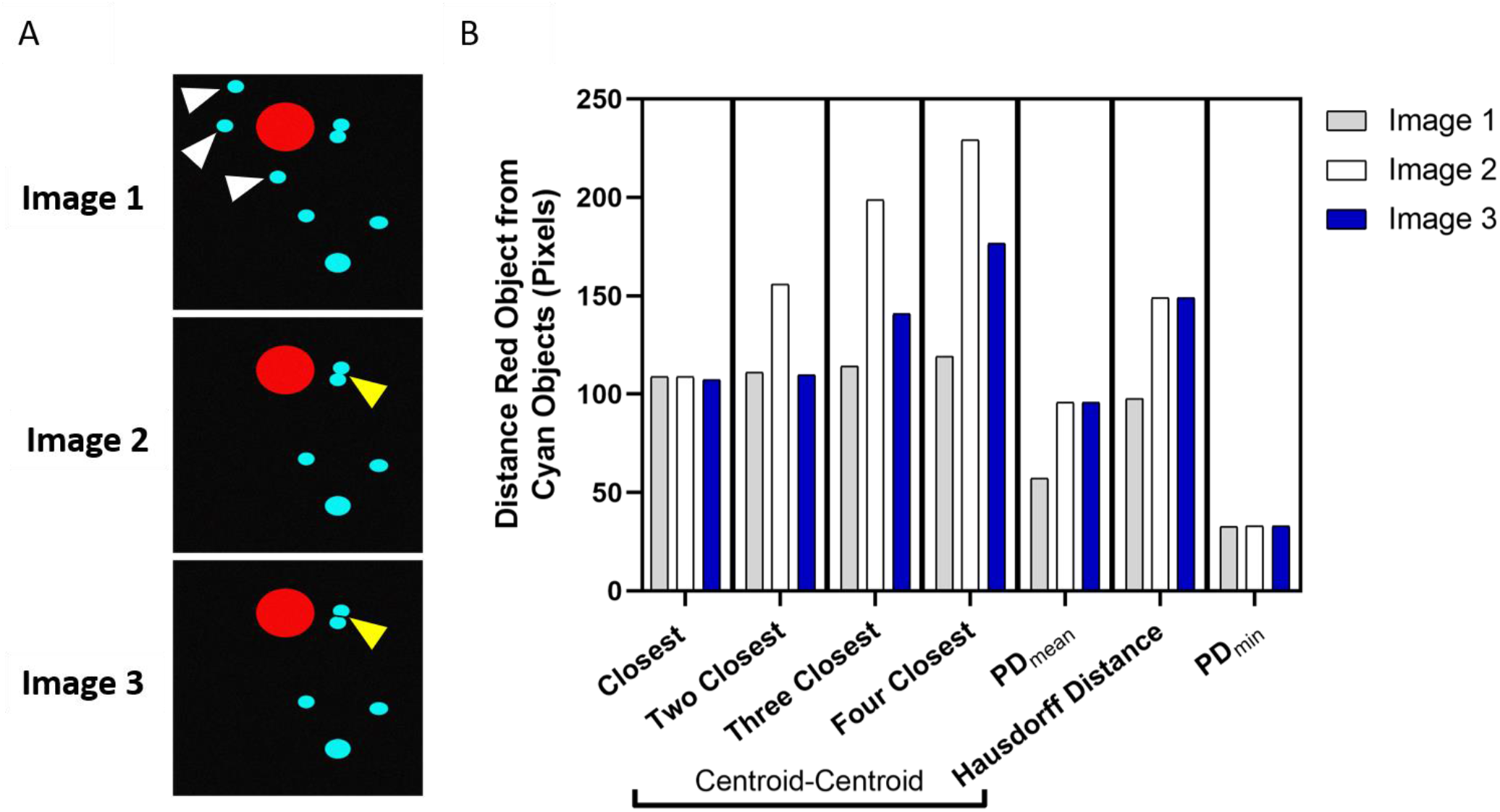
PD_mean_ values provide more robust descriptions of biomarker proximity then centroid-centroid measurements. (A) Images generated using Inkscape software with random pixel noise added in Python to better resemble true microscopy images. The red signal in each image is in the same spatial location each time. Image 2 is a duplicate of Image 1 with the cyan objects indicated by white arrow heads removed. Image 3 is a duplicate of Image 2 with a small region of null signal (indicated by the yellow arrow heads) added to divide the object, into two objects. (B) Distance measurements of each image. Centroid-centroid distances were calculated in DiAna using default parameters ^7^. The ‘Closest’ centroid-centroid measurement describes the centroid-centroid distance between the red object and the nearest blue object. The ‘Two Closest’ – ‘Four Closest’ centroid-centroid measurements describe the average (mean) between the red object and the two – four closest cyan objects. PD_mean_, Hausdorff Distance and PD_min_ measurements were calculated using PyBioProx as previously described.

There is a 2.18 fold increase in the PD_mean_ distance of the average (mean) *S. aureus* cluster to LAMP-1 (1.5 hr, 0.11 ± 0.011 µm; 24 hr, 0.24 ± 0.013 µm (SEM)) at the 24 hr time point compared to the 1.5 hr time point. These PD_mean_ values are displayed in Fig 4B as a frequency distribution. Differences in the percentage distribution of bacterial PD_mean_’s can be observed at the early and late time points. While the percentage distributions of both 1.5 hr and 24 hr time points are both positively skewed, this effect is far more pronounced at 1.5 hrs. Indeed, at 1.5 hrs, 58.0 % of *S. aureus* clusters have a PD_mean_ of ≤ 0.05 µm, which drops to 19.5 % at 24 hrs. Therefore, in this infection model, PD_mean_ measurements corroborate the results of the manual analysis (Fig 3B) suggesting that *S. aureus* MSSA476 is less likely to be found encapsulated by LAMP-1 at late compared to early time points. This is consistent with previous findings that *S. aureus* is capable of escape from LAMP-1 positive vesicles ^17,22^.

Increases in *S. aureus-*LAMP-1 distances are also observed between early and late timepoints using the PD_min_ measurement (1.5 hr, 0.00 ± 0.00 µm; 24 hr, 0.035 ± 0.0071 µm (SEM)) (Fig 4E). However, this dataset highlights the lack of resolving power of the PD_min_, as 100 % and 83.32 % of bacteria at the 1.5 hr and 24 hr timepoints (respectively) had a PD_min_ of 0.00 µm (Fig 4F). A small increase in Hausdorff Distances was observed (1.5 hr, 0.53 ± 0.032 µm; 24 hr, 0.68 ± 0.023 µm (SEM)) (Fig 4C). However, there is no clear distinction between the frequency distributions of the 1.5 hr and 24 hr time points (Fig 4D).

While the Hausdorff Distance performed well in a 2D ‘computer-generated’ dataset (Fig 2D) it was not able to convincingly identify true differences in biomarker proximity in a 3D ‘real-world’ dataset (Fig 4D). The poor performance of the Hausdorff and PD_min_ measurements likely reflects the inherent issues of extrema-based distance measurements. A single ‘erroneous’ pixel in the segmentation process would be sufficient to throw either of these measurements off. For example, if a strongly LAMP-1 associated *S. aureus* object had a *single* perimeter pixel erroneously detected in a Z-slice where it was not ‘truly’ present, the Hausdorff Distance could be erroneously inflated. The PD_mean_ is likely to be more robust to such ‘noise’ as the statistic reflects the average of *all* PD measurements. Collectively, this data demonstrates that the PD_mean_ measurement (but not the Hausdorff Distance or PD_min_ measurements) is capable of identifying known differences in biomarker proximity in a real-world dataset.

### 3. Advantages of PD_mean_ measurements over paired-object measurements

The PD_mean_ measurement describes the proximity of a fluorescent object relative to *all* surrounding fluorescent objects in another channel. Previously described distance-based analysis methods instead calculate the distance between *pairs* of fluorescent objects, e.g. by measuring the distances between the centroids of paired-objects ^5,7,10,11^. The ‘DiAna’ ImageJ plug-in, implements a version of this paired-object approach ^7^.

#### Advantage 1: PD_mean_ measurements are less reliant on analyst decision-making to accurately describe biomarker proximity

To illustrate the advantages of PD_mean_ measurements over the paired-object approach, we generated artificial images of a red ‘biomarker’ surrounded by cyan ‘biomarkers’ (Fig 5A). ‘Image 1’ contains seven cyan objects; four clustered tightly to the red object and three located further away. ‘Image 2’ is a duplicate of ‘Image 1’ but with three of the four closest cyan objects (indicated by white arrowheads) removed. The red object in ‘Image 1’, therefore, has an increased proximity to cyan objects when compared to the red object in ‘Image 2’. To define which object-object pairing to collect measurements from, the DiAna ImageJ plug-in first ranks them by distance. The user then determines the number of measurements to collect. For example, in ‘Image 1’, the options range from collecting only the closest (red-cyan) measurement to collecting the seven closest measurements. While a histogram showing all possible object-object distances *does* describe the relationship of the red object to the cyan objects; generating a histogram for each object-of-interest in a microscopy image results in unwieldy data, especially in large microscopy datasets with many objects-per-image. A single value describing the colocalisation of each object-of-interest (e.g., an average of centroid-centroid distances) is likely to be more usable.

The number of centroid-centroid measurements to incorporate into this single value is not necessarily obvious and profoundly influences the outcome of the analysis. For example, only when averaging two or more centroid-centroid pairings, is the red object in ‘Image 1’ described as having increased proximity to cyan objects then the red object in ‘Image 2’ (Fig 5B). The appropriate number of measurements to take will likely vary from image to image. Taking both too few or too many paired measurements may result in poor detection of real differences in the proximity of biomarkers. Either, by excluding relevant measurements from the calculation or, by diluting the signal with irrelevant measurements. By contrast, PD_mean_ and Hausdorff distance measurements both show apparent differences between ‘Images 1 and 2’ with no further analysis decisions needed.

#### Advantage 2: PD_mean_ measurements reduce errors associated with an overreliance on accurate object detection

Making centroid-centroid measurements requires, by definition, for object detection to be performed upon all fluorescent channels that are to be analysed. PD measurements, by contrast, require object detection in only one (red) channel, while the second (cyan) channel is merely thresholded. This reduced dependency on accurately detecting objects limits the impact of small changes in object detection on the average spatial colocalisation of an image. For example, in Fig 5A, ‘Image 3’ is a duplicate of ‘Image 2’ with a small region of null signal (indicated by the yellow arrowheads) added to divide the closest cyan object such that two objects are now detected. Because of this small change in the image, when averaging the two, three and four closest centroid-centroid measurements, the red object in Image 3 is incorrectly described as being 29.6 %, 29.1 % and 22.3 % (respectively) more spatially associated with cyan objects then the red object in ‘Image 2’ (Fig 5B). Indeed, if considering only one or two centroid-centroid measurements, ‘Image 3’ appears slightly more spatially colocalised than ‘Image 1’ despite the reduced number of cyan objects in close proximity to the red object.

The Hausdorff distance measurement identifies no difference between ‘Image 2’ and ‘Image 3’ while the PD_mean_ measurement correctly identifies a tiny (0.02 pixel) increase in the spatial proximity of the red object in ‘Image 3’ compared to ‘Image 2’, reflecting the small region of null signal added in ‘Image 3’. The PD_min_ measurement could not correctly identify differences between any of the sample images. Collectively, this shows that within this dataset, of the trialled measurements, PD_mean_ was most able to accurately identify changes in spatial proximity.

## Conclusions

In this paper, we have defined the PD_mean_, a simple measurement that describes the relative proximity of one set of fluorescent biomarkers to another. We have also introduced PyBioProx: an open-source image-analysis tool that quantifies the relative proximity of fluorescent biomarkers in 2D and 3D microscopy datasets. Despite being an *object-based* analysis method, we found that the PD_mean_ measurement was able to accurately identify changes in colocalisation in simulated data meant for assessing *pixel-based* colocalisation metrics (Fig 2B & E). We then assessed the capacity of this analysis to make biological relevant observations.

Using the PD_mean_ measurement, we identified significant differences in the spatial proximity of intracellular *S. aureus* to a marker of an intracellular compartment (LAMP-1) at early and late time points (Fig 4A & B), confirming a manual analysis (Fig 3B). We also illustrated instances where currently used paired centroid-centroid measurements would be poorly able to accurately describe the proximity of one set of biomarkers to another (Fig 5A). PD_mean_ measurements were able to overcome these shortcomings (Fig 5B). Collectively these results suggest that PyBioProx and PD_mean_ measurements can function as a powerful means of analysing the relative spatial proximity of fluorescent biomarkers.

## References

1. Verveer, P. J. & Bastiaens, P. I. H. Quantitative microscopy and systems biology: seeing the whole picture. Histochem. Cell Biol. 130, 833–843 (2008).

2. Bolte, S. & Cordelières, F. P. A guided tour into subcellular colocalization analysis in light microscopy. J. Microsc. 224, 213–232 (2006).

3. Li, Q. et al. A syntaxin 1, Gαo, and N-type calcium channel complex at a presynaptic nerve terminal: analysis by quantitative immunocolocalization. J. Neurosci. 24, 4070–4081 (2004).

4. Manders, E. M. M., Stap, J., Brakenhoff, G. J., Driel, R. V. & Aten, J. A. Dynamics of three-dimensional replication patterns during the S-phase, analysed by double labelling of DNA and confocal microscopy. J. Cell Sci. 103, 857–862 (1992).

5. Obara, B., Jabeen, A., Fernandez, N. & Laissue, P. P. A novel method for quantified, superresolved, three-dimensional colocalisation of isotropic, fluorescent particles. Histochem. Cell Biol. 139, 391–402 (2013).

6. Lachmanovich, E. et al. Co-localization analysis of complex formation among membrane proteins by computerized fluorescence microscopy: application to immunofluorescence co-patching studies. J. Microsc. 212, 122–131 (2003).

7. Gilles, J.-F., Dos Santos, M., Boudier, T., Bolte, S. & Heck, N. DiAna, an ImageJ tool for object-based 3D co-localization and distance analysis. Methods 115, 55–64 (2017).

8. Jaskolski, F., Mulle, C. & Manzoni, O. J. An automated method to quantify and visualize colocalized fluorescent signals. J. Neurosci. Methods 146, 42–49 (2005).

9. Moser, B., Hochreiter, B., Herbst, R. & Schmid, J. A. Fluorescence colocalization microscopy analysis can be improved by combining object-recognition with pixel-intensity-correlation. Biotechnol. J. 12, 1600332 (2017).

10. Nelson, C. J. et al. Blobs and curves: object-based colocalisation for plant cells. Funct. Plant Biol. 42, 471 (2015).

11. Sassmann, S. et al. An immune-responsive cytoskeletal-plasma membrane feedback loop in plants. Curr. Biol. 28, 2136-2144.e7 (2018).

12. Walt, S. van der et al. scikit-image: image processing in Python. PeerJ 2, e453 (2014).

13. Otsu, N. A threshold selection method from gray-level histograms. IEEE Trans. Syst. Man Cybern. 9, 62–66 (1979).

14. Zinchuk, V., Wu, Y. & Grossenbacher-Zinchuk, O. Bridging the gap between qualitative and quantitative colocalization results in fluorescence microscopy studies. Sci. Rep. 3, 1365 (2013).

15. Flannagan, R., Heit, B. & Heinrichs, D. Antimicrobial mechanisms of macrophages and the immune evasion strategies of Staphylococcus aureus. Pathogens 4, 826–868 (2015).

16. Kinchen, J. M. & Ravichandran, K. S. Phagosome maturation: going through the acid test. Nat. Rev. Mol. Cell Biol. 9, 781–795 (2008).

17. Flannagan, R. S., Heit, B. & Heinrichs, D. E. Intracellular replication of Staphylococcus aureus in mature phagolysosomes in macrophages precedes host cell death, and bacterial escape and dissemination. Cell. Microbiol. 18, 514–535 (2016).

18. Jubrail, J. et al. Inability to sustain intraphagolysosomal killing of Staphylococcus aureus predisposes to bacterial persistence in macrophages. Cell. Microbiol. 18, 80–96 (2016).

19. Surewaard, B. G. J. et al. Identification and treatment of the Staphylococcus aureus reservoir in vivo. J. Exp. Med. 213, 1141–1151 (2016).

20. Custódio, R. et al. Characterization of secreted sphingosine-1-phosphate lyases required for virulence and intracellular survival of Burkholderia pseudomallei. Mol. Microbiol. 102, 1004–1019 (2016).

21. Dallenga, T. et al. M. tuberculosis-induced necrosis of infected neutrophils promotes bacterial growth following phagocytosis by macrophages. Cell Host Microbe 22, 519-530.e3 (2017).

22. Grosz, M. et al. Cytoplasmic replication of Staphylococcus aureus upon phagosomal escape triggered by phenol-soluble modulin α. Cell. Microbiol. 16, 451–465 (2014).

23. Kubica, M. et al. A potential new pathway for Staphylococcus aureus dissemination: The silent survival of S. aureus phagocytosed by human monocyte-derived macrophages. PLoS ONE 3, e1409 (2008).

24. Kim, S. H. & Allebach, J. P. Optimal unsharp mask for image sharpening and noise removal. J. Electron. Imaging 14, 023005 (2005).

