## Supplementary Figures 1 & 2 for "A Quantitative Measurement to Describe the Relative Proximity of Fluorescent Biomarkers" (from another version of this preprint: version 1, 2020-12-09)

### Supplemental Figures

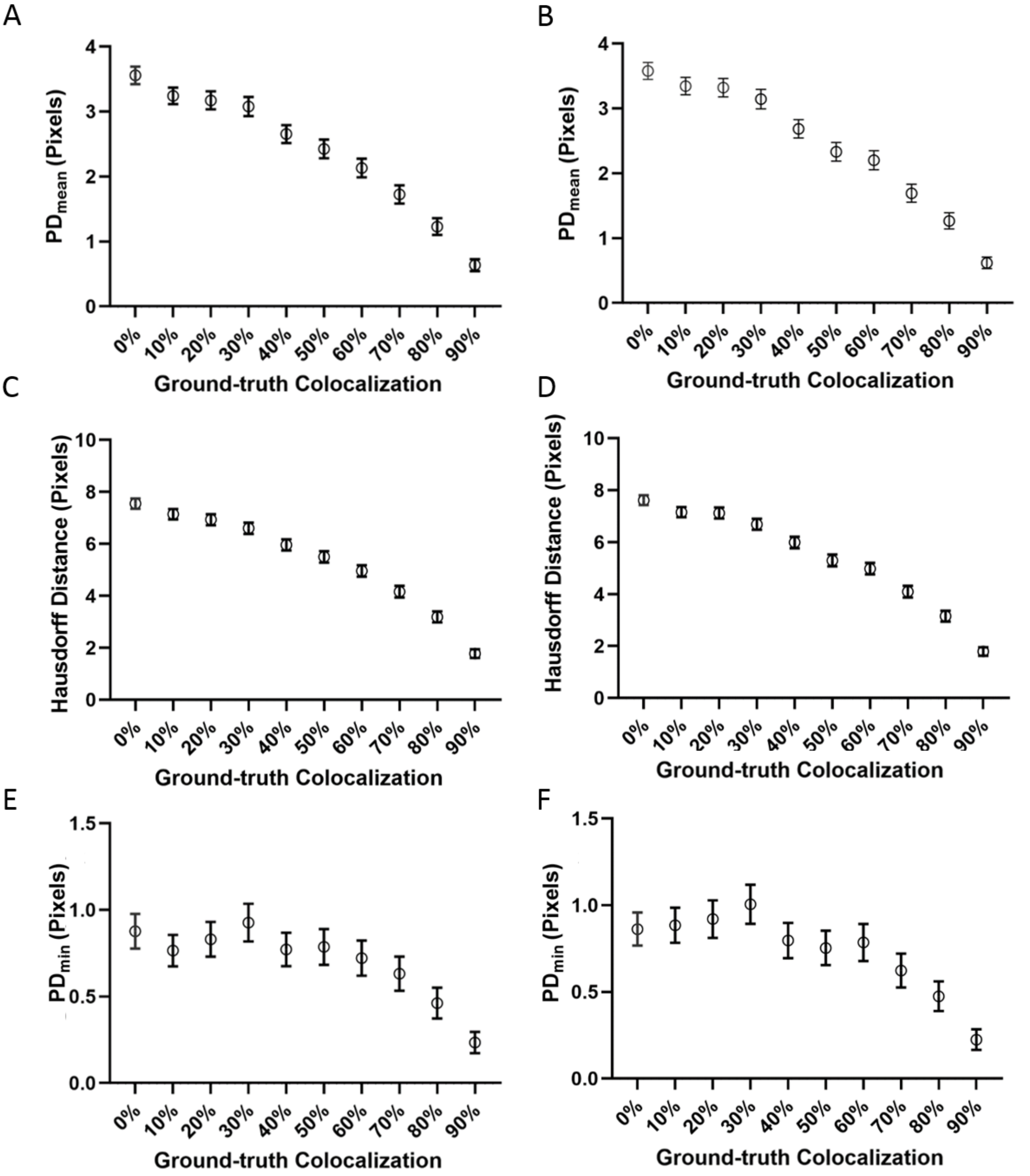


**Supp. Fig 1 – $PD_{mean}$, Hausdorff Distance but not $PD_{min}$ accurately identify ground-truth colocalisation in CBS datasets 2 and 3.** $PD_{mean}$ (A-B), Hausdorff Distance (C-D) and $PD_{min}$ measurements describing the relative proximity of objects CBS "Image Set 2" (A, C & E) and CBS "Image Set 3" (B, D & F). Distances are from Red objects to Blue objects in "Image Set 2" and from Green objects to Blue objects in "Image Set 3". Values represent the mean distance of a Red/Green object from Blue objects in each image ± 95 % CI.

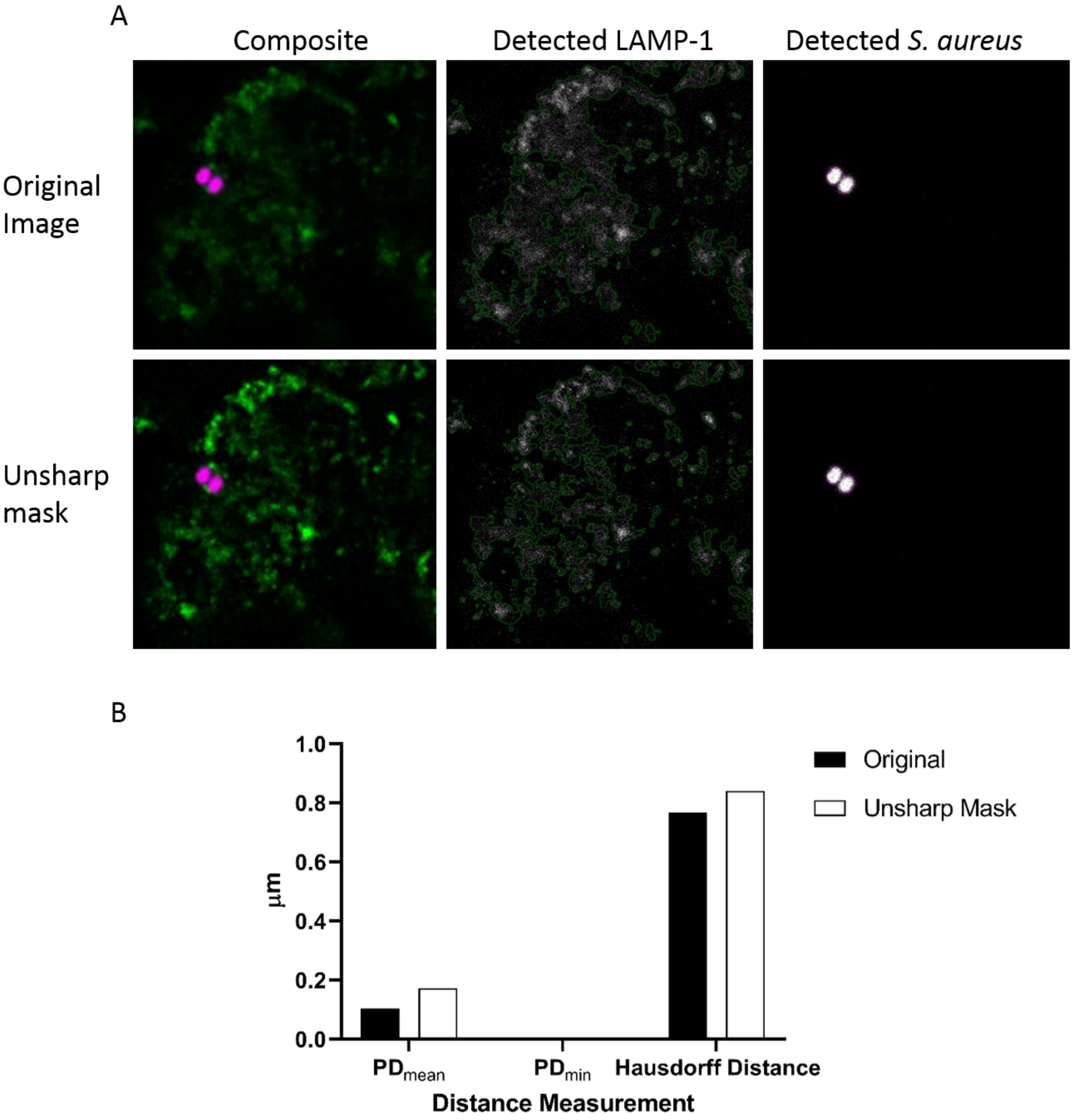


**Supp. Fig 2 - Impact of Unsharp Mask Pre-processing.** (A) The objects identified by PyBioProx in an image pre-processed using unsharp masking (Unsharp mask) and without (Original) unsharp masking in the LAMP-1 channel only. Both images were pre-processed with a gaussian blur (sigma of 1). (B) The $PD_{mean}$, $PD_{min}$ and Hausdorff Distance measurements of the images shown in (A).
